# FiloQuant reveals increased filopodia density during DCIS progression

**DOI:** 10.1101/125047

**Authors:** Guillaume Jacquemet, Ilkka Paatero, Alexandre F. Carisey, Artur Padzik, Jordan S. Orange, Hellyeh Hamidi, Johanna Ivaska

**Affiliations:** Turku Centre for Biotechnology, University of Turku, FIN-20520, Finland.; Center for Human Immunobiology, Texas Children’s Hospital, Houston, TX 77030, USA; Department of Pediatrics, Pathology and Immunology, Baylor College of Medicine, Houston, TX 77030, USA; Department of Biochemistry, University of Turku, FIN-20520, Finland.

## Abstract

Filopodia are commonly observed cellular protrusions in vitro and in vivo. Defective filopodia formation is linked to several pathologies including cancer, wherein actively protruding filopodia, at the invasive front, and filopodia-mediated probing of the microenvironment accompanies cancer cell dissemination. Despite wide biological significance, delineating the function of these finger-like protrusions in more complex systems remains technically challenging, particularly hindered by lack of compatible methods to quantify filopodia properties. Here, we present FiloQuant, a freely available ImageJ plugin, to detect filopodia and filopodia-like protrusions in both fixed and live-cell microscopy data. We demonstrate that FiloQuant can extract quantifiable information including protrusion dynamics, density and length from multiple cell types and in a range of microenvironments, such as during collective or single cancer cell migration in 2D and 3D, in fixed neuronal cultures, in activated natural killer cells and in sprouting endothelial cells in vivo. In cellular models of breast ductal carcinoma in situ (DCIS) we reveal a link between filopodia formation at the cell-matrix interface, during collective invasion and in 3D tumour spheroids, with the previously reported local invasive potential of these breast cancer models in vivo. Finally, using intravital microscopy, we observed that tumour spheroids display prominent filopodia in vivo, supporting a potential role for these protrusions during tumorigenesis.

## Introduction

The extension of membrane protrusions is a prominent morphological feature during many cellular processes and serves as an important mechanism to probe the extracellular matrix (ECM) and to ascertain the appropriate cellular response. Cellular protrusions are broadly classified in function of membrane shape and/or size and primarily include lamellipodia, membrane blebs, filopodia and filopodia-like protrusions (Chhabra and Higgs, 2007; Petrie and Yamada, 2012). Filopodia are thin, finger-like projections exploited widely by different cell types, including neurons, endothelial cells, epithelial cells, fibroblasts and immune cells (Heckman and Plummer, 2013; Mattila and Lappalainen, 2008; Jacquemet et al., 2015), wherein they contribute to cellular communication (Sagar et al., 2015), directional cell migration (Jacquemet et al., 2015) and the establishment of cellcell junctions (Biswas and Zaidel-Bar). For instance, in neurons, filopodia have been implicated in neurite outgrowth and in the formation of both neurite and dendritic spines (Mattila and Lappalainen, 2008; Björkblom et al., 2012). In vivo, filopodia have been reported to contribute to processes such as endothelial sprouting and angiogenesis (Wakayama et al., 2015; Phng et al., 2013), ECM deposition and remodeling (Sato et al., 2017), epithelial sheet migration during wound healing and dorsal closure (Wood et al., 2002; Millard and Martin, 2008) and embryonic development (Fierro-González et al., 2013).

Filopodia may also contribute to pathological conditions including cancer and brain disorders (Kanjhan et al., 2016; Jacquemet et al., 2015). We and others have reported that filopodia and filopodia-like protrusions are extensively used by cancer cells to support directional single cell migration and invasion as well as survival at distant metastatic sites (Jacquemet et al., 2013a; Paul et al., 2015; Arjonen et al., 2014; Jacquemet et al., 2016; Shibue et al., 2013, 2012). In addition, the expression of a number of filopodia regulatory proteins has been shown to correlate with poor patient survival in multiple cancer types, the downregulation of which impedes cancer metastasis in animal models (Arjonen et al., 2014; Li et al., 2014; Yap et al., 2009). Therefore, targeting filopodia formation could prove a viable strategy to impair cancer cell metastasis (Jacquemet et al., 2016). However, cancer cell dissemination is an intricate multi-step process (Gupta and Massagué, 2006) and the significance of filopodia at every stage of the metastatic cascade is not clear.

In spite of their wide biological importance, filopodia remain poorly studied primarily due to technical difficulties. In particular, filopodia are difficult to observe, especially in vivo, owing to their small size, the absence of specific markers and due to an often labile nature particularly affected by fixation protocols (Wood and Martin, 2002; Sato et al., 2017). In addition, automatic quantification of filopodia properties remains a challenge, despite the availability of dedicated tools, and therefore filopodia features are often described using manual analyses. To our knowledge, currently available tools to quantify filopodia include Filodetect (Nilufar et al., 2013), CellGeo (Tsygankov et al., 2014) and ADAPT (Barry et al., 2015), each with unique strengths and shortcomings (Table 1). Limitations of these tools include requirement for proprietary software (i.e., MATLAB and MATLAB Image Processing Toolbox), lack of customizable options to improve filopodia detection, selective dedication to live-cell data or to fixed samples only, designation for single cells only, quantification of filopodia numbers but not density and the usage of an unmodifiable and/or complex code source that precludes addition of extra functionalities by nonexperts.

**Table 1:** Comparison of FiloQuant with previously described filopodia analysis software.

|  | **FiloDetect**<br>(Nilufar et al., 2013) | **CellGeo**<br>(Tsygankov et al., 2014) | **ADAPT**<br>(Barry et al., 2015) | **FiloQuant** (this study) |
| --- | --- | --- | --- | --- |
| Requirement for proprietary software | Yes (MATLAB) | Yes (MATLAB) | No (ImageJ) | No (ImageJ) |
| Analysis of single images | Yes | No | No | Yes |
| Analysis of movies / filopodia tracking | No | Yes | Yes | Yes in combination with a freely available ImageJ plugin |
| Optimized for single cells | Yes | Yes | Yes | Yes |
| Optimized for cell sheets | No | No | No | Yes |
| Quantification of filopodia length and number | Yes | Yes | Yes | Yes |
| Quantification of edge length to measure filopodia density | No | Yes (indirectly) | No | Yes |
| Options to improve faint filopodia detection | No | No | No | Yes |
| Option to filter filopodia in function of size | No | Yes | Yes | Yes |
| Option to filter filopodia in function of proximity to cell/colony edge | No | No | No | Yes |
| Batch mode available | No | Yes | Yes | Yes |
| Customizable code (easily) | No | No | No | Yes |
| Easy access to update | No | No | Yes | Yes |
| Analysis of lamellipodia | No | Yes | Yes | No |
| and/or blebs |  |  |  |  |

In the process of addressing these limitations, we developed FiloQuant to detect filopodia and filopodia-like protrusions and to extract quantifiable data on protrusion dynamics, density and length from both fixed and live-cell microscopy images. Here, we provide three versions of FiloQuant (software 1: single image analysis; software 2: semi-automated; software 3: fully automated for large analyses and/or live-cell imaging), on the freely available ImageJ platform, each designed with a different purpose in mind and with alternative levels of user control over the analysis. Using FiloQuant, we demonstrate that filopodia can be successfully detected in different cells and in a range of microenvironments including during collective or single cancer cell migration in 2D and 3D, in fixed neuronal cultures and in sprouting endothelial cells in vivo, regardless of the imaging modality. Using FiloQuant to analyse a cellular model of breast cancer progression we report that filopodia density and length, during collective cell invasion and in 3D spheroids, correlates with previously reported local invasive potential through the basement membrane in vivo (Miller et al., 2000; Behbod et al., 2009; Lodillinsky et al., 2016). Finally using intravital microscopy, we report that tumour spheroids in vivo display a high number of filopodia, potentially highlighting an important role for filopodia in regulating tumorigenesis.

## Results

### FiloQuant, an ImageJ tool to rapidly quantify filopodia length and density under different cellular contexts

We developed FiloQuant as a plugin for the ImageJ software with inter operating systems compatibility (Schindelin et al., 2012). In brief, FiloQuant works by first defining the cell/colony edge in an input image following intensity based thresholding and removal of long thin protrusions such as filopodia from the plasma membrane (Figure 1A). In parallel, the same input image is separately enhanced to optimize filopodia detection (Figure 1B) and is then superimposed on the filopodia-erased cell-edge image to specifically isolate filopodia at the cell boundary. The number and length of these cell-edge filopodia are then automatically analysed by FiloQuant. Filopodia density can also be determined by calculating the ratio of filopodia number to edge length (extracted by FiloQuant from the edge detection image) (Figure 1A).

**Figure 1:**
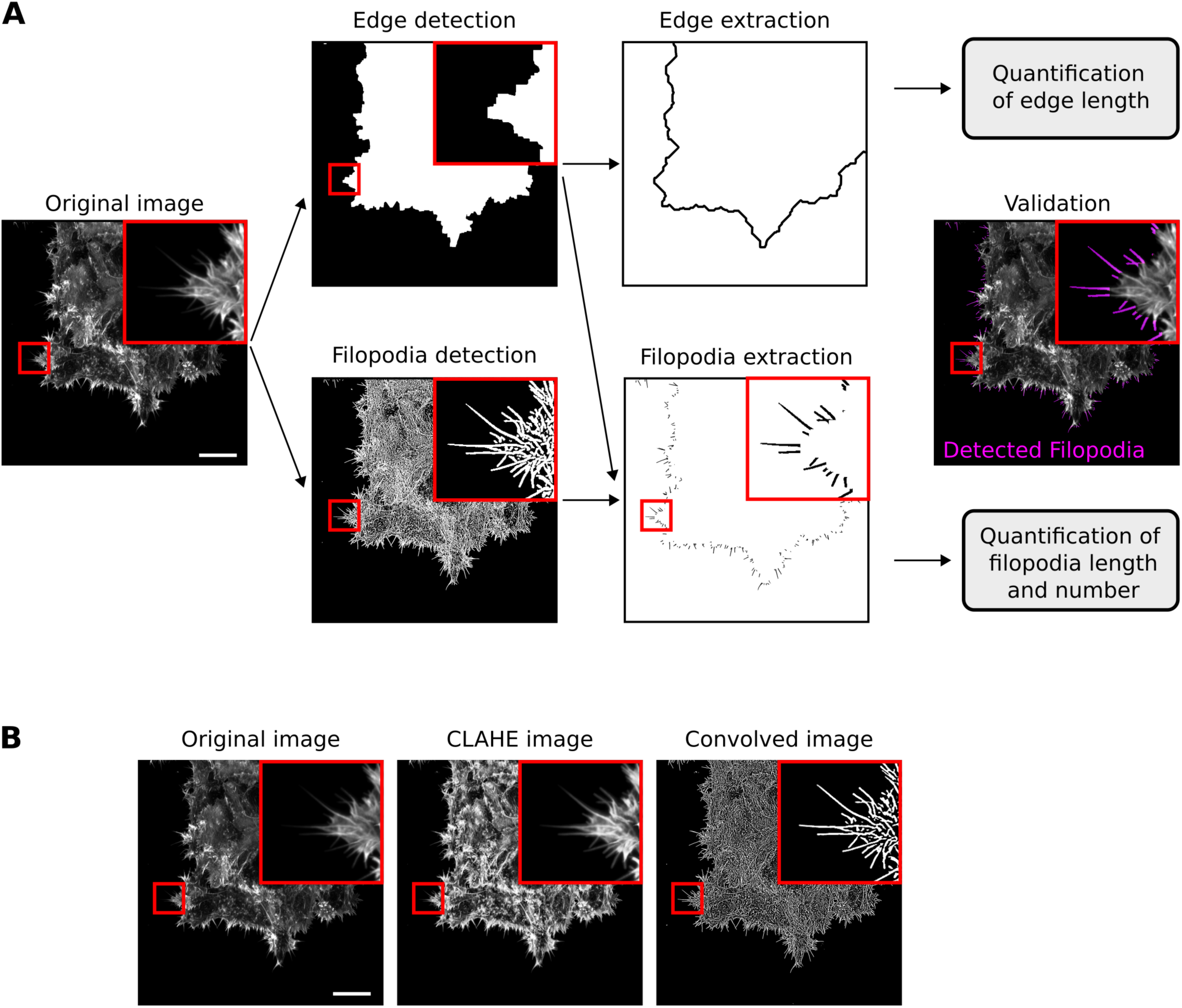
FiloQuant, an ImageJ tool to rapidly quantify filopodia length and density. **A:** Work-flow depicting FiloQuant analysis of filopodia density and length. Representative images obtained at the different stages of analysis are displayed. Briefly, the original image (input) undergoes two parallel processing steps. In the top panel, the cell edge is defined and detected by intensity based thresholding and by “erasing” the filopodia (edge detection). In the bottom panel, the image is enhanced to optimize detection of faint filopodia without introducing noise (filopodia detection). The resulting images are subtracted to isolate edge filopodia (filopodia extraction) and filopodia number and length are automatically analysed. Detected filopodia are highlighted in magenta in the final image. Filopodia density can be also quantified by determining the ratio of filopodia number to edge length (extracted from the edge detection image). The original image shows MCF10 ductal carcinoma in situ (DCIS.COM) cells invading collectively through a fibrillar collagen gel (circular invasion assay), stained for actin and imaged using a spinning disk confocal (SDC) microscope (100x objective, Orca Flash 4 camera, scale bar = 20 μm). **B:** Images illustrate how two of the settings (CLAHE and Convolve) available in FiloQuant can help to improve the detection of faint filopodia (scale bar = 20 μm).

To make this software as easy to use as possible, FiloQuant (single image analysis version; software 1) contains step-by-step user validation of the various processing stages to help users achieve optimal settings for filopodia detection. This is especially important as efficient detection of filopodia can vary from image to image, even when acquired under similar settings, mainly due to the small size of these structures resulting in weak signals. Several enhancements as well as filtering options are available in FiloQuant to improve filopodia detection and we provide here a detailed manual explaining how to use FiloQuant (supplementary information). In particular, among the enhancement strategies tested, a fairly conservative convolution kernel (Figure 1B) was found to be very effective in improving detection of faint filopodia.

We demonstrate that FiloQuant can be broadly applied to successfully detect filopodia from an assortment of cell images acquired on different microscopes. These include cells migrating collectively (Figure 2A) or as single cells (Figure 2B-C) in various environments such as on 2D fibronectin (Figure 2B) or on 3D cell-derived matrices (Figure 2C). In addition, we show that FiloQuant can detect filopodia in neurons, which have a more complex morphology, (Figure 2D) and can distinguish filopodia-like protrusions in activated natural killer cells (Figure 2E).

**Figure 2:**
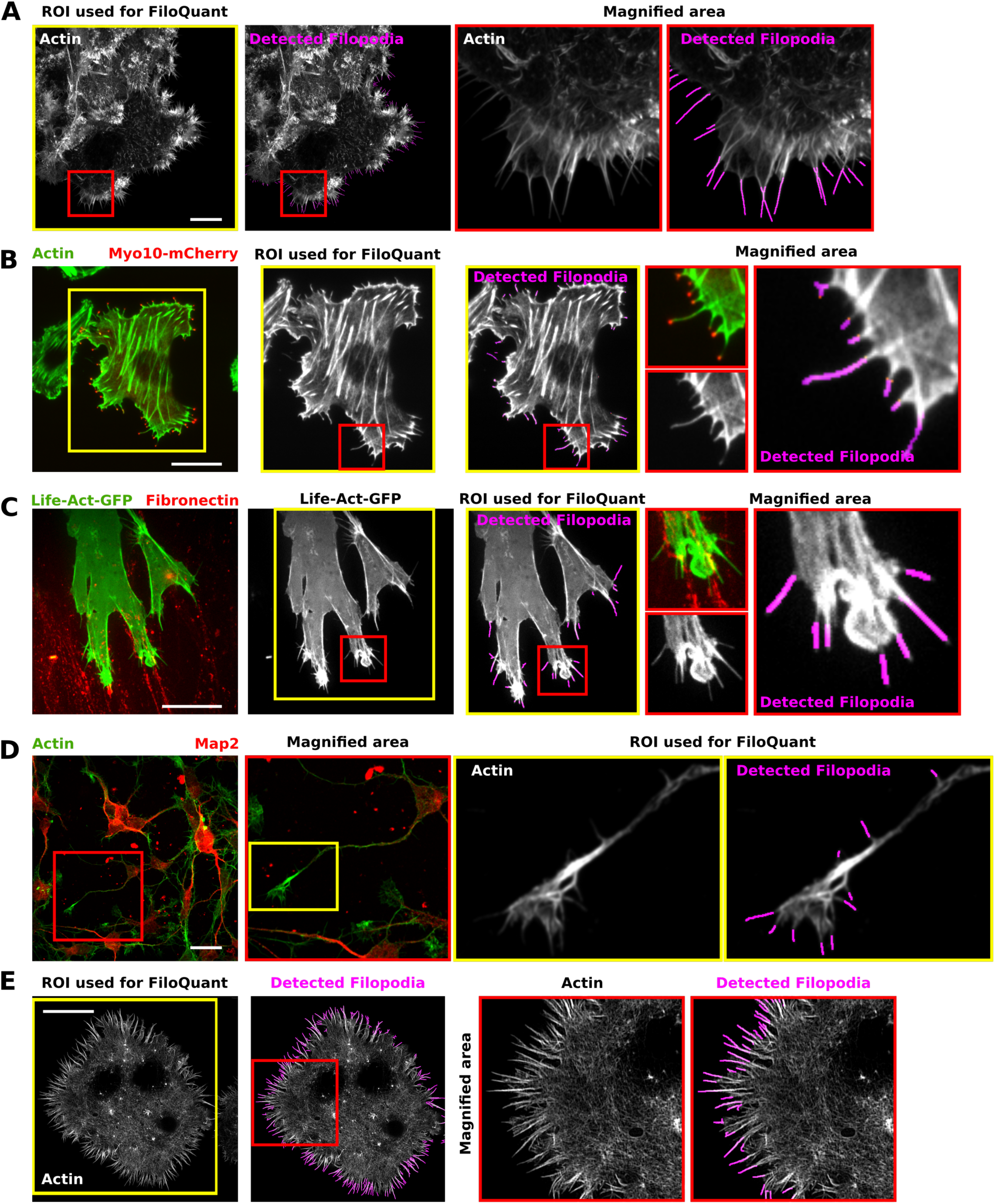
FiloQuant can be used to detect filopodia under different cellular contexts and imaging modalities. **A-E:** To illustrate the versatility of FiloQuant, filopodia detection was evaluated in images acquired from different cell types and using different imaging modalities. The single image analysis version of FiloQuant (Software 1) was used in all cases. DCIS.COM cells migrating collectively were stained for actin and imaged using an SDC microscope (100x, Orca Flash 4 camera, scale bar = 20 μm) (A). MDA-MB-231 cells transiently expressing mCherry-Myosin-X (to visualise filopodia tips) were plated for 2 h on fibronectin (FN), stained for actin and imaged using a total internal reflection fluorescence (TIRF) microscope (scale bar = 20 μm) (B). A2780 cells transiently expressing mEmerald-Lifeact and migrating on cell-derived matrices (CDMs) in the presence of exogenous FN (labelled with alexa Fluor 568) were imaged live on an SDC microscope (63x oil objective, Evolve 512 EMCCD camera, scale bar = 20 μm) (C). Primary rat hippocampal neurons plated on laminin were fixed, stained for actin and MAP2 (neuronal marker) and imaged using an SDC microscope (100x objective, Orca Flash 4 camera, scale bar = 20 μm) (D). (E) NK-92 natural killer cells were seeded on antibody-coated glass (anti-CD18 and anti-NKp30) for 20 minutes before being fixed, stained for actin and imaged by TIRF-SIM. The actin-rich filopodia-like protrusions were then detected using FiloQuant (scale bar = 10 μm). For all panels, red insets denote magnified area and yellow insets denote the region of interest (ROI) analysed by FiloQuant. Filopodial protrusions detected by FiloQuant are displayed in magenta.

### Detection of filopodia in vivo, during angiogenesis, using intravital imaging and FiloQuant

To further test the flexibility of FiloQuant, we analysed filopodia properties in vivo in images acquired using high resolution intravital microscopy (Figure 3). In living organisms, filopodia are employed by multiple cell types (Jacquemet et al., 2015) including endothelial tip cells, which generate long filopodial protrusions during the process of angiogenesis (Wakayama et al., 2015). To visualize angiogenesis in vivo, we treated genetically modified zebrafish embryos expressing GFP in the endothelium (genotype Tg(kdrl:GFP); roy -/-; mitfa -/-) with a low concentration of latrunculin B (Figure 3) previously reported to inhibit filopodia formation in endothelial tip cells (Phng et al., 2013). The sprouting segmental arteries of the embryos were then imaged live on a spinning disk confocal microscope and z projections were created for analysis (Figure 3A).

**Figure 3:**
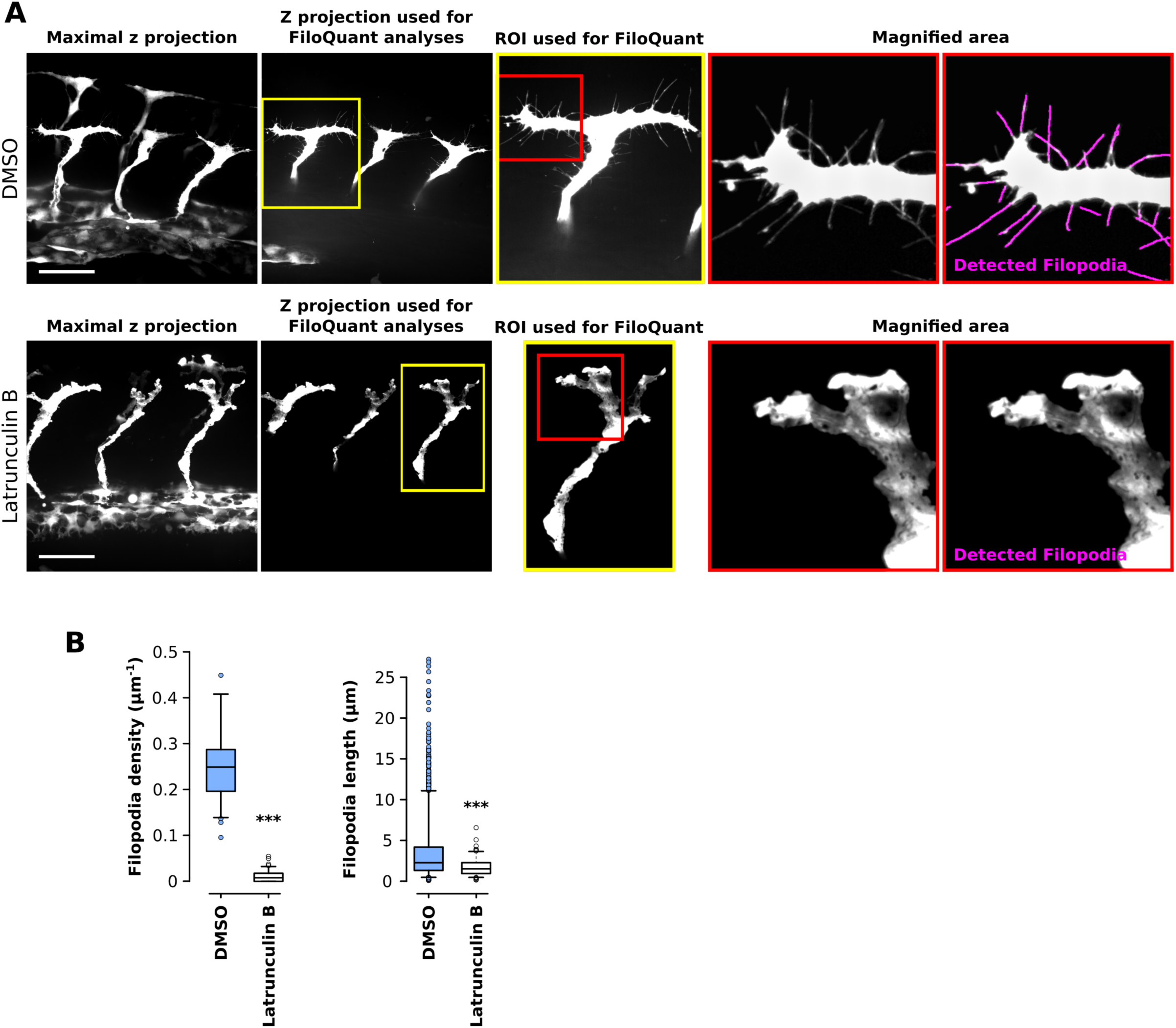
Filopodia can be detected and quantified in sprouting endothelia in the developing zebrafish embryo using intravital imaging and FiloQuant. **A-B:** Transgenic zebrafish embryos expressing GFP in the endothelium (genotype Tg(kdrl:GFP); roy-/-; mitfa-/-) were treated with DMSO (1%) or latrunculin B (150 ng/ml) from 25 hours postfertilization (hpf) to 29 hpf. The embryos were then anesthetized and mounted in low-melting point agarose on glass-bottom dishes. Z stack images of the sprouting segmental arteries were obtained live using an SDC microscope (long working distance 63x water objective, Orca Flash 4 camera). Representative maximal z projection and the z projection used for FiloQuant analyses are shown (A). In addition, ROI (yellow square) used for FiloQuant analyses, magnified area (red squares), and filopodia detected using FiloQuant (magenta) are displayed (scale bar = 20 μm). Quantification of filopodia density and filopodia length using the semi-automated version of FiloQuant are displayed as Tukey box plots (B) (DMSO, 15 embryos imaged; latrunculin B, 17 embryos imaged; Filopodia density: DMSO, 60 ROI analyzed; latrunculin B, 71 ROI analyzed; Filopodia length: DMSO, 2232 filopodia measured; latrunculin B, 138 filopodia measured; ***P value < 5.49×10^-25^). The Tukey box plots represent the median and the 25th and 75th percentiles (interquartile range); points are displayed as outliers if 1.5 times above or below the interquartile range; outliers are represented by dots. Statistical analysis: Student’s t-test (unpaired, two-tailed, unequal variance).

To analyze larger datasets while retaining control over the settings used to analyze each image, and to modify these settings on the fly to improve the accuracy of detection, a second version of FiloQuant was created (semi-automated; software 2). Using the semi-automated version of FiloQuant, we were able to rapidly and efficiently detect and analyze filopodia properties in sprouting endothelial cells, and, as previously reported (Phng et al., 2013), demonstrated a significant inhibition in filopodia density following latrunculin B treatment (Figure 3B). Moreover, the length of the remaining filopodia was also substantially reduced compared to those in the control DMSO treated embryos. This example, in addition to those presented in Figure 2, clearly demonstrates that FiloQuant is a powerful tool to detect and quantify filopodial features under different cellular contexts and imaging modalities.

### Filopodia density and length in vitro correlate with increased malignancy in ductal carcinoma in situ

Filopodia and filopodia-like protrusions are prominent features of migrating/invading cancer cells in vitro (Jacquemet et al., 2013a; Paul et al., 2015; Petrie and Yamada, 2012). However the significance of filopodia at the different stages of the metastatic cascade remains unclear. In breast cancer, metastasis is initiated by cells breaking through a basement membrane to escape the tumour in situ and to invade locally in the surrounding stroma. To study local invasion, a cellular model of breast cancer that recapitulates the different stages of ductal carcinoma in situ (DCIS) progression in animal models and importantly mimics the human disease (Lodillinsky et al., 2016; Behbod et al., 2009; Miller et al., 2000) was used. This model is composed of three cell lines: i) normal immortalized breast epithelial cells (MCF10A), ii) premalignant H-Ras transformed MCF10A variants (MCF10AT cell line) that are tumorigenic as xenografts, and iii) tumorigenic and invasive MCF10A variants (MCF10DCIS.COM cell line) (Miller et al., 2000; Dawson et al., 1996). When growing MCF10A, MCF10AT and MCF10DCIS.COM (DCIS.COM) cells under growth-factor-reduced (GFR) Matrigel, the actin protrusions formed at the invading edge of cell colonies were strikingly different (Figure 4A). In particular, the highly invasive DCIS.COM cells exhibited substantially more filopodia compared to MCF10A and MCF10AT cells (Figure 4A). To further validate this observation, MCF10A and DCIS.COM cells were plated in cell culture inserts, and left to invade for three days through an overlay of fibrillar collagen or GFR Matrigel or left in media only (Figure S1; Figure 4B). Filopodia density and length at the invasive edge were then quantified using FiloQuant (Figure 4C-E). Importantly, DCIS.COM colonies displayed higher filopodia density, in addition to longer filopodia, compared to MCF10A colonies regardless of the composition of the microenvironment (Figure 4B, 4C and 4E). Surprisingly, the composition of the microenvironment did impact on filopodia density within each cell population (Figure 4B and 4D-E). In particular, in DCIS.COM cells, invasion into GFR Matrigel induced very dense arrays of short filopodia compared to media alone or fibrillar collagen. Invasion into fibrillar collagen, in turn, triggered higher filopodia density compared to media alone without affecting filopodia length. Similarly, in MCF10A cells, invasion through GFR Matrigel and fibrillar collagen increased filopodia density, but not filopodia length, compared to media alone (Figure 4B and 4D-E). Taken together, these data are indicative of a prominent role for the cell microenvironment in regulating filopodia formation, previously believed to be dictated primarily by cancer cell intrinsic properties such as oncogenes.

**Figure 4:**
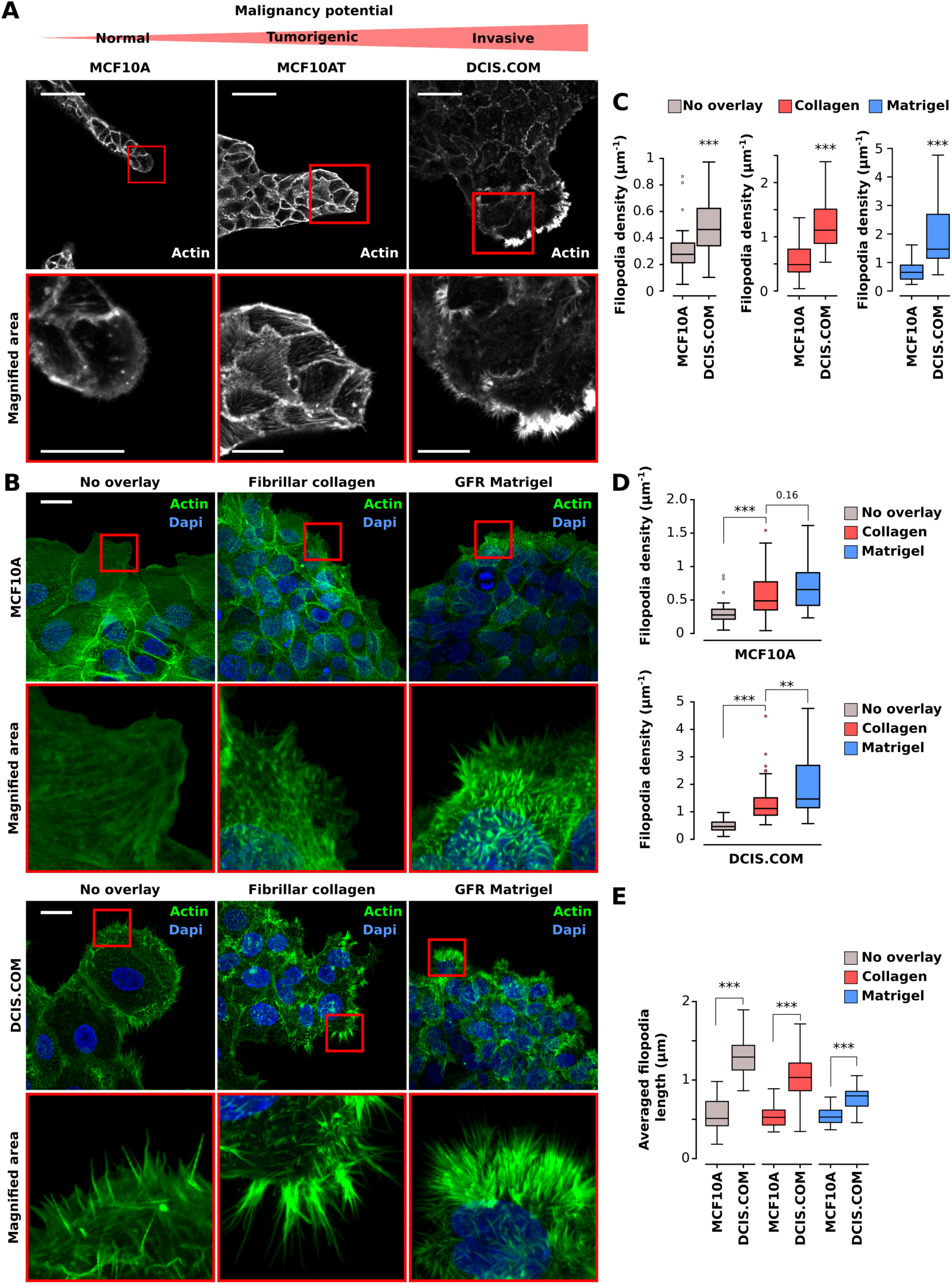
Filopodia density and length in circular invasion assays correlate with reported degree of cancer cell malignancy. **A:** MCF10A, MCF10AT and DCIS.COM cells were left to migrate into GFR Matrigel for 14 days, fixed, stained and imaged using a confocal microscope (scale bar = 50 μm; magnified area, scale bar = 20 μm). **B-E:** MCF10A and DCIS.COM cells were plated in circular invasion assays and left to invade through GFR Matrigel, fibrillar collagen I or media (no overlay) for 3 days. Cells were then fixed, stained for actin and Dapi, and imaged using an SDC microscope (100x objective, Orca Flash 4 camera). For each condition, representative maximal z projection and magnified area (red squares) are displayed (scale bar = 20 μm). Filopodia density was compared between cells (C, ***P value <1.54×10^-4^) and in the same cells in different overlay conditions (D, **P value = 0.002, ***P value < 9.99×10^-05^). Average filopodia length was also calculated in each cell line (E, ***P value < 3.46×10^-10^). Results from three independent experiments are displayed as Tukey box plots (condition, fields of view analysed; MCF10A no overlay, 43; MCF10A fibrillar collagen I, 31; MCF10A GFR Matrigel, 37; DCIS.COM no overlay, 30; DCIS.COM fibrillar collagen I, 73; DCIS.COM GFR Matrigel, 37). Statistical analysis: Student’s t-test (unpaired, two-tailed, unequal variance).

### Analysis of filopodia dynamics using FiloQuant

As filopodia formation appeared to be different in MCF10A and DCIS.COM cells (figure 4), we set out to further characterize these structures using live-cell imaging. To visualize actin dynamics in live cells, MCF10A and DCIS.COM cell lines expressing Life-act-RFP (Life-act MCF10A and Life-act DCIS.COM, respectively) were generated using lentivirus (see methods). Importantly, constitutive expression of Life-act-RFP did not perturb filopodia density, filopodia length, or proliferation in DCIS.COM cells (Figure 5A-B). To study filopodia dynamics, Life-act MCF10A and Life-act DCIS.COM cells were plated in circular invasion assays (Figure S1) and left to invade for three days through fibrillar collagen or GFR Matrigel before being imaged live using a spinning disk confocal microscope (Figure 5C and Video 1). Notably, DCIS.COM cells were able to invade collectively and efficiently through fibrillar collagen and GFR Matrigel, whereas MCF10A cells migrated mostly within the cell sheet and did not invade (Figure 5C and Video 1).

**Figure 5:**
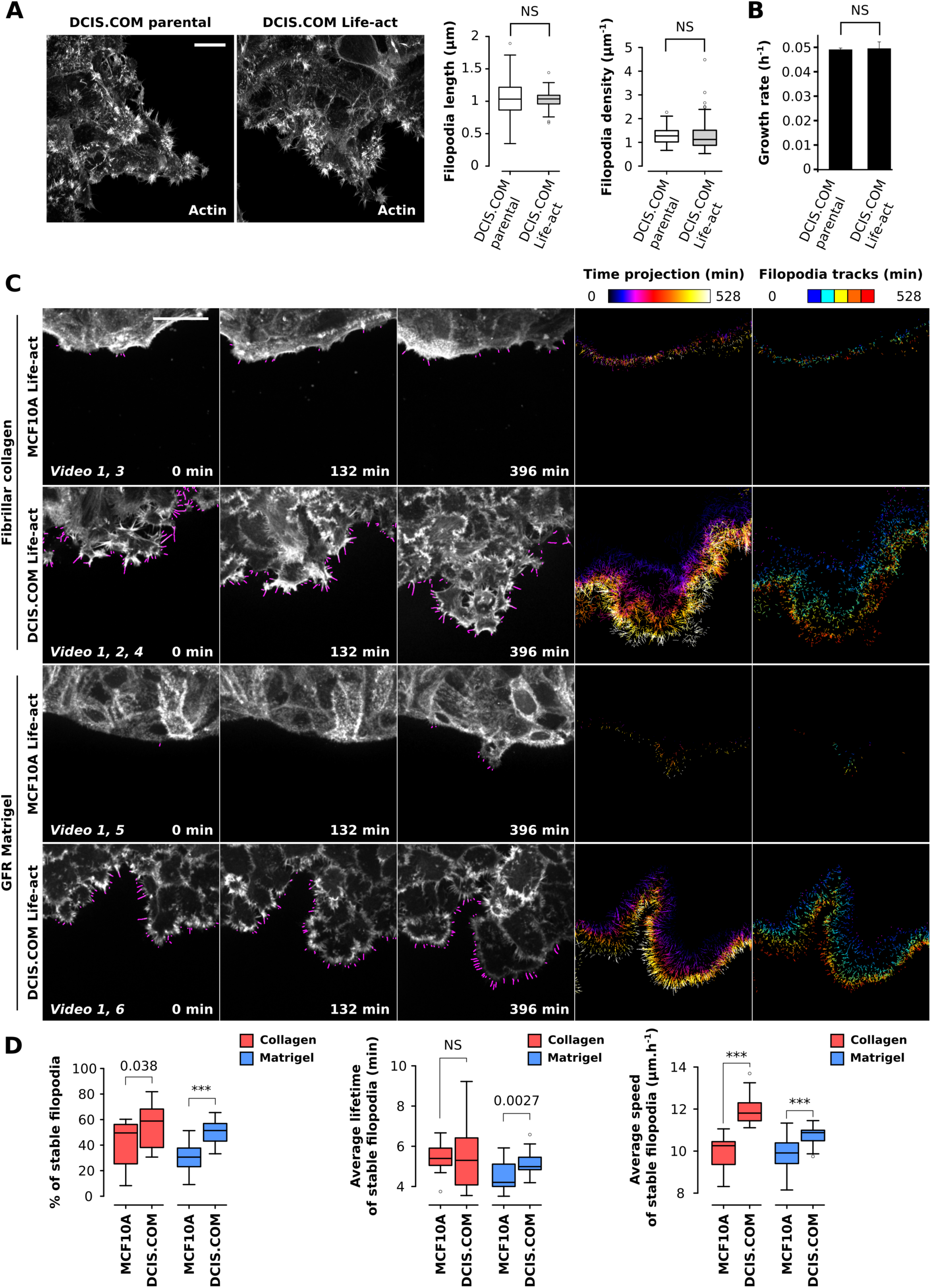
Filopodia dynamics can be analysed using FiloQuant in circular invasion assays. **A-B:** To analyse filopodia dynamics, DCIS.COM cells stably expressing Life-act mRFP (DCIS.COM Life-act) were generated and validated by comparison to parental DCIS.COM cells. Cells were plated in circular invasion assays and left to invade through fibrillar collagen I for 3 days. Cells were then fixed, stained for actin, and imaged using an SDC microscope (100x objective, Orca Flash 4 camera). Representative maximal z projection (scale bar = 20 μm), filopodia density, as well, as filopodia length measured using FiloQuant are displayed. Results from three independent experiments are displayed as Tukey box plots (condition, fields of view analysed; DCIS.COM, 73; DCIS.COM Life-act, 41) (A). In addition, rate of cell growth was recorded using an IncuCyte ZOOM^®^ live-cell microscopy incubator and the confluency method. Results are from three independent experiments (B). **C-D:** MCF10A and DCIS.COM cells stably expressing Life-act-mRFP were plated in circular invasion assays and left to invade through fibrillar collagen I or GFR Matrigel for 3 days before being imaged live using an SDC microscope (100x objective, Evolve 512 EMCCD camera) for over 9 h (1 picture every 3 min). Filopodia were then automatically detected using the automated version of FiloQuant and tracked using TrackMate (ImageJ based tracking tool). For each condition, stills of single z planes with filopodia detected by FiloQuant (magenta) are displayed (scale bar = 25 μm) at different time points. In addition, time projection of detected filopodia as well as filopodia tracks are shown. The time projections were directly generated by FiloQuant and are colour coded as a function of time. For filopodia tracking, tracking files, generated by FiloQuant, were entered into TrackMate for automated tracking. Filopodia tracks generated by TrackMate are colour coded as a function of the track starting time (track index) (C). For each condition, the percentage of stable filopodia (lifetime > 3 min), the average lifetime of stable filopodia and the average speed of stable filopodia were quantified from three independent experiments and displayed as Tukey box plots (condition, movies analysed; MCF10A Life-act fibrillar collagen I, 14; MCF10A Life-act GFR Matrigel, 19; DCIS.COM Life-act fibrillar collagen I, 23; DCIS.COM Life-act GFR Matrigel, 30; ***P value < 2.79×10^-04^) (D). Statistical analysis: Student’s t-test (unpaired, two-tailed, unequal variance).

For analysis of large datasets, where the same settings can be applied to many images at once at the beginning of the analysis, a third version of FiloQuant was created (fully automated; software 3). This is especially useful for high-throughput assays and/or to analyse filopodia properties and dynamics from live-cell imaging data. Movies acquired from Life-act MCF10AT and Life-act DCIS.COM cells were analyzed using this automated version of FiloQuant (software 3) and the “stack analysis” option (see FiloQuant guide provided as supplementary information) for sequential processing of images. In particular, this mode enables the analysis of filopodia properties per frame, creates a time projection of all detected filopodia (Figure 5C and Videos 2-6) and provides a tracking file (Videos 2 and 7), which can then be used to automatically track filopodia dynamics using freely available ImageJ tools such as TrackMate (Tinevez et al., 2017). Using FiloQuant and TrackMate, we found that DCIS.COM cells generate a higher proportion of stable filopodia (> 3 min lifetime) compared to MCF10A cells regardless of the composition of the microenvironment (Figure 5D). In addition, filopodia generated by DCIS.COM cells moved more rapidly in the direction of invasion compared to filopodia generated by MCF10A cells. This observation is likely to be linked to the ability of the DCIS.COM cells to invade through the microenvironment.

### Filopodia density in 3D spheroids correlates with increased malignancy

Within a 3D environment, MCF10A and MCF10A-derived cell lines form spheroids encapsulated within a basement membrane that is believed to better recapitulate the in vivo situation. In particular, MCF10A spheroids establish and maintain a polarity axis that results in the formation of a lumen by apoptotic clearance, while MCF10AT and DCIS.COM spheroids fail to polarize or to form a lumen, the absence of which is considered to be a hallmark of tumorigenesis (Hernandez-Fernaud et al., 2017; Debnath and Brugge, 2005; Imbalzano et al., 2009). To investigate whether these MCF10A cell-line-derived spheroids also exhibit distinct filopodia similar to those observed in 2D (Figure 4 and 5), we plated single MCF10A, MCF10AT and DCIS.COM cells on GFR Matrigel and monitored filopodia formation in spheroids at different time points post-plating using a confocal microscope (Figure 6A) (Debnath et al., 2003). Strikingly, DCIS.COM spheroids displayed very prominent edge filopodia at day 7 and 14 post-plating compared to MCF10A or MCF10AT spheroids (Figure 6A). These dense arrays of filopodia persisted in the larger DCIS.COM spheroids at day 21 post-plating (Figure 6B). We analysed filopodia density in MCF10A and DCIS.COM spheroids at day 5 (using single z planes), when the entire spheroid was still contained within the imaging window at high magnification and resolution, using a spinning disk confocal microscope (Figure 6C). Quantification of filopodia density, using FiloQuant, clearly demonstrated extensive filopodia assembly in DCIS.COM cells at the spheroid edge, whereas in MCF10A spheroids, filopodia were largely absent from the borders (Figure 6C).

**Figure 6:**
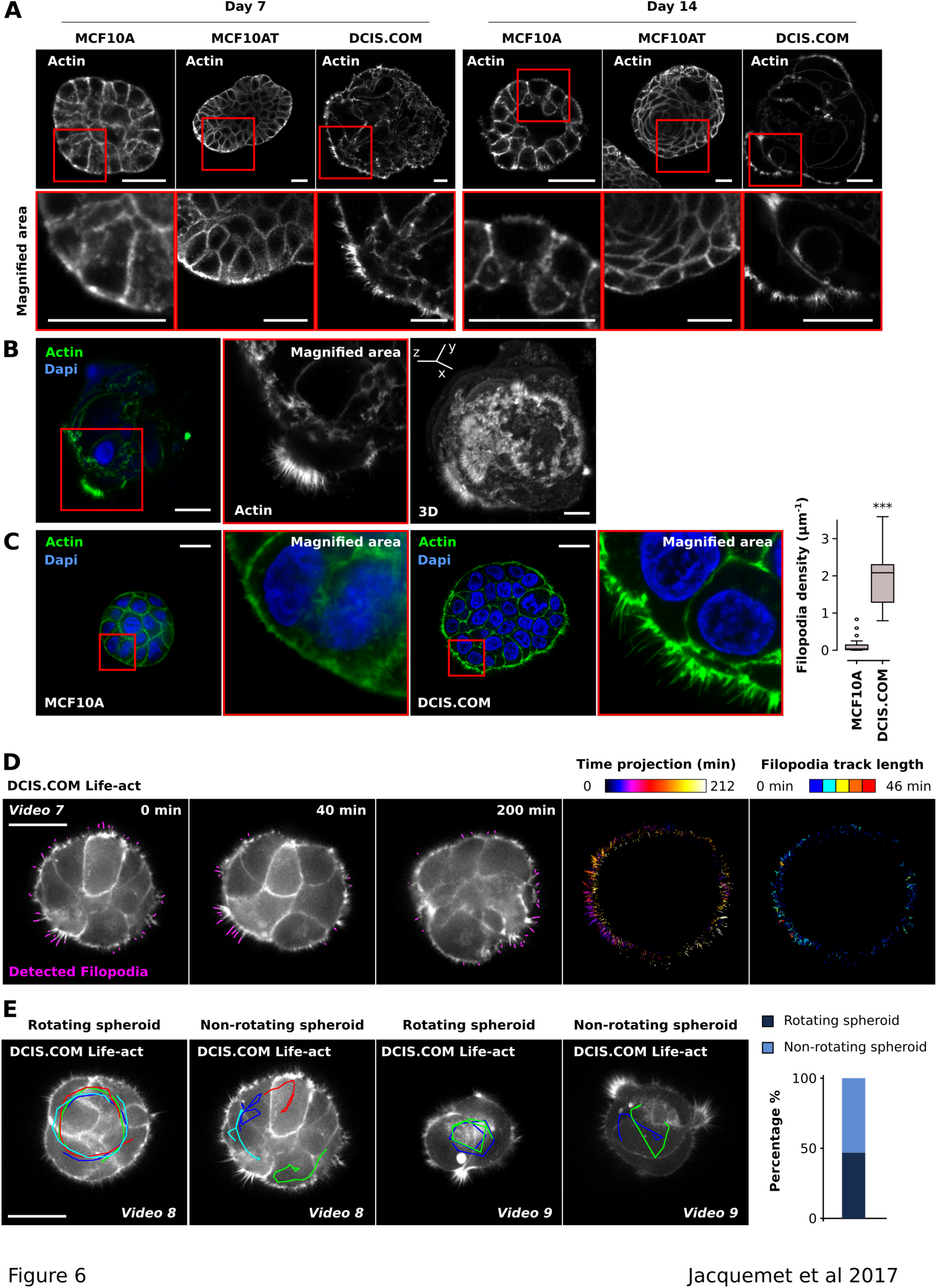
Filopodia density in 3D spheroids correlates with increased malignancy. **A:** MCF10A, MCF10AT and DCIS.COM cells were seeded as single cells on GFR Matrigel and left to form spheroids over 7 or 14 days, fixed, stained and imaged using a confocal microscope. For each condition, representative single z planes as well as magnified areas (red squares) are displayed (scale bars = 25 μm). **B:** DCIS.COM cells were seeded as in A and left to form spheroids for 21 days, fixed, stained and imaged using a confocal microscope. A representative single z plane, magnified region (red square) and an Imaris-generated 3D reconstruction are displayed (scale bars = 25 μm). **C:** MCF10A and DCIS.COM cells were seeded as in A and allowed to form spheroids for 5 days, fixed, stained and imaged using an SDC microscope (100x objective, Orca Flash 4 camera). Representative single z planes and magnified areas (red square) are displayed (scale bars = 20 μm). Filopodia density at spheroid borders was quantified using FiloQuant. Results from two independent experiments are displayed as Tukey box plots (condition, fields of view analysed; MCF10A, 18; DCIS.COM, 11; ***P value = 3.06 × 10^-05^). **D-E:** DCIS.COM Life-act cells were seeded as in A and allowed to form spheroids for 3 days before being imaged live using an SDC microscope (100x objective, Evolve 512 EMCCD camera) for over 3 h (1 picture every 2 min). Filopodia were then automatically detected using FiloQuant and tracked using TrackMate as in figure 5C. Stills of single z planes with filopodia detected by FiloQuant (magenta), a time projection of the detected filopodia and filopodia tracks are displayed (scale bar = 25 μm). The time projections are color coded as a function of time. Filopodia tracks are colour coded as a function of the track length to highlight filopodia stability (D). To highlight cellular behaviour within spheroids, migration tracks (red, light and dark blue and green) are displayed on top of four different spheroids. In addition, the proportion of rotating spheroids over non-rotating spheroids is displayed. Results are from three independent experiments (81 movies analyzed) (E).

To investigate filopodia dynamics in 3D, Life-act DCIS.COM cells were plated as single cells on GFR Matrigel and allowed to form spheroids for 3 days before being imaged live using a spinning disk confocal microscope (Videos 7-9). Filopodia were then automatically detected and tracked using FiloQuant and TrackMate (Figure 6D and Video 7). These analyses revealed that, in 3D, DCIS.COM cells can produce extremely stable filopodia with a long lifetime approaching 40-50 min (Figure 6D). Surprisingly, in contradiction with previous reports (Tanner et al., 2012; Wang et al., 2013) we found that cancer-cell-line-derived spheroids can undergo rotational movement as demonstrated by approximately 50% of DCIS.COM-generated spheroids (Figure 6E).

### Tumour spheroids generate filopodia in vivo

We next sought to investigate whether DCIS.COM filopodia observed in vitro are present in vivo. As filopodia in vivo are often lost upon chemical fixation (Wood and Martin, 2002; Sato et al., 2017), we used intravital microscopy to directly visualise filopodia in an in vivo setting. Specifically, Life-act DCIS.COM cells were injected into the pericardial cavity of zebrafish embryos and imaged live 24 h post-injection using a spinning disk confocal microscope (Figure 7). DCIS.COM cells were able to survive in the pericardial cavity and to form 3D spheroids similar to those observed in 3D GFR Matrigel (Figure 7A and B). High resolution intravital imaging of these tumours revealed the presence of dense filopodial networks at the spheroid border (Figure 7B-D and Videos 10-11), thus, confirming that filopodia formation is not limited to in vitro cultures and that filopodia are generated in vivo by the invasive DCIS.COM breast cancer cells.

**Figure 7:**
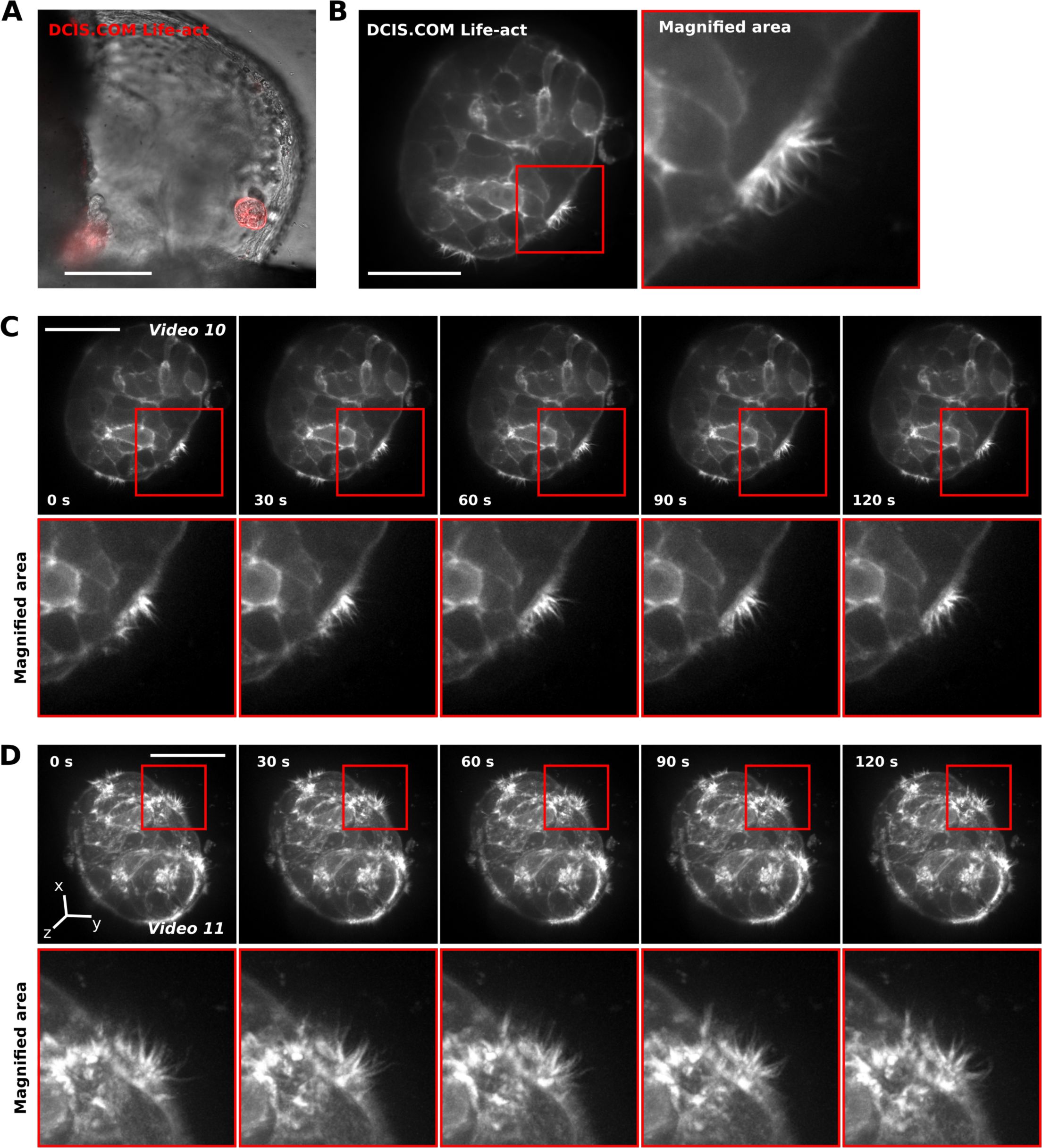
Tumour spheroids generate filopodia in vivo. **A-B:** DCIS.COM Life-act cells were injected into the pericardial cavity of zebrafish embryos and imaged live using an SDC microscope 24 h post injection. Prior to imaging, the embryos were anesthetized and mounted in low-melting point agarose on glass-bottom dishes. A representative image of a DCIS.COM Life-act spheroid growing in the zebrafish embryo pericardial cavity (20x objective, Orca Flash 4 camera, scale bar = 100 μm) (A) and a single z plane (63x water objective, Orca Flash 4 camera, scale bar = 40 μm) and magnification (B) are shown. **C-D:** Live imaging of the DCIS.COM Life-act spheroid, shown in B, using an SDC microscope (63x water objective, Evolve 512 EMCCD camera, 1 frame per 10s). Single z planes (C, scale bar = 20 μm) and Imaris-generated 3D reconstructions (D, scale bar = 20 μm) are displayed. Red insets denote magnified areas.

Altogether, we developed FiloQuant, a novel software that can be successfully applied in different settings to detect and quantify filopodia properties. Using FiloQuant, we focused on investigating the presence and dynamics of filopodia in cell models reflecting different stages of cancer progression; however we expect that FiloQuant would also be useful in quantifying filopodia in other contexts and/or to quantify other finger-like protrusions such as retraction fibers or nanotubes. In addition, despite the broad functions offered by the software, we appreciate that other users may require extra functionalities or the ability to include FiloQuant within larger analysis routines. To this end and to facilitate easy modification, the simple ImageJ macro language used to generate FiloQuant has been fully annotated (software 1-3) and can therefore be edited with limited coding knowledge. For easy installation and rapid distribution, we have also deposited FiloQuant on an ImageJ Update site (see methods).

## Discussion

The extensive employment of filopodia in vivo by different cell types and mounting in vitro and clinical evidence linking filopodia and/or filopodia regulatory pathways to disease (Jacquemet et al., 2015), emphasize the need for more comprehensive analyses of filopodia function. Here, we developed FiloQuant, an ImageJ-based computational tool, in order to simplify quantification of filopodia and applied this tool to investigate filopodial properties in cell models of cancer progression.

In cancer, filopodium-like protrusions have been demonstrated to enable the outgrowth of macrometastases and cancer survival at distal sites (Shibue et al., 2013, 2012). Moreover, we and others have shown a dense arrangement of actin-rich filopodial structures at the invasive front of cancer cells in vitro that is required for single cell invasion in a 3D setting (Jacquemet et al., 2013a; Paul et al., 2015; Arjonen et al., 2014; Jacquemet et al., 2016). Here, using FiloQuant and a cellular model of breast cancer that recapitulates the different stages of DCIS (Miller et al., 2000; Behbod et al., 2009; Dawson et al., 1996), we identified an association between the formation of stable and dense arrays of filopodia, in tumour spheroids and during collective invasion, with the previously reported potential of these breast cancer cell models to pierce through the basement membrane in vivo (Behbod et al., 2009; Lodillinsky et al., 2016). These data suggest that filopodia may also be important for collective local invasion and the initiation of metastasis; however filopodia requirement for collective cell invasion requires further investigation. Intriguingly, in our study, breast cancer cell invasion through a gel composed of basement membrane constituents induced the highest filopodia density. This observation supports the notion that filopodia formation is not merely a cancer-cell intrinsic property but is strongly reinforced by the surrounding matrix potentially via integrin receptor signalling (Jacquemet et al., 2016).

Despite the existence of convincing in vitro and clinical evidence, the use of filopodia by cancer cells in vivo remains poorly documented. Using intravital microscopy, we report that tumour cells injected into the pericardial cavity of zebrafish embryos form spheroids in vivo and that these spheroids display a high number of filopodia. The exact function of these filopodia in this situation is not clear; however, filopodia analyses in appropriate animal models mimicking the different steps of the metastatic cascade, combined with advances in high resolution microscopy, are likely to help unravel the role of these filopodia in cancer progression in vivo.

In addition, we report here that DCIS.COM spheroids can display rotational movement in 3D Matrigel. Rotational movement of spheroids was previously reported in several non-malignant mammary epithelial cells and associated with their ability to polarize (Tanner et al., 2012) and to assemble a basement membrane (Wang et al., 2013). Surprisingly, these previous studies described a loss of rotational motion in several cancer cell lines including MCF-7, T47D, Caco-2 and Panc-1 cells and described this phenomenon as a hallmark of a malignant phenotype (Tanner et al., 2012; Wang et al., 2013). Our observation that rotational movement is not fully lost in DCIS.COM spheroids is now challenging this previous conclusion and suggests that lack of rotational movement may not always correlate with malignancy.

Filopodia are important for many other pathophysiological processes in addition to cancer and we show that FiloQuant can be applied to quantify filopodia properties (length, density and dynamics) across different cell types, microenvironments and image acquisition techniques and therefore presents several advantages over previously described Filopodia analysis tools (Nilufar et al., 2013; Barry et al., 2015; Tsygankov et al., 2014) (Table 1): i) to the best of our knowledge, this is the only currently available software capable of extracting filopodia properties from either single images (individual or in batch) or live-cell imaging data, ii) FiloQuant can be applied to detect edge protrusions in both single cells and in cell colonies, iii) FiloQuant analyses include quantification of filopodia density, iv) FiloQuant contains modifiable parameters, in addition to filtering options based on filopodia size and proximity to the cell or colony edge, that improve the detection of faint filopodia. The only systems requirement, the installation of the ImageJ/Fiji (Schindelin et al., 2012; Schneider et al., 2012) platform (with the addition of several plug-ins already packaged in Fiji), enables easy and free dissemination of the software throughout the cell biology community. Importantly, FiloQuant’s basic ImageJ macro language can be easily modified by non-experts to add extra functionalities and/or as a means to incorporate FiloQuant in larger analysis routines. Extra functionalities could include quantification of protein recruitment to filopodia or evaluation of other filopodia and/or cell edge parameters such as filopodia angles or straightness and cell shape. In addition, FiloQuant outputs can be effortlessly coupled to existing ImageJ plug-ins. As an example we connected FiloQuant to Trackmate (Tinevez et al., 2017) to automatically track filopodia dynamics. A detailed user guide to FiloQuant is provided here as supplementary information. Furthermore, to allow customization to address specific needs, the original FiloQuant source code has been fully annotated and deposited in an open online repository (ImageJ update site).

FiloQuant was designed specifically to assess filopodia properties; however, we anticipate a broader application for FiloQuant in the analysis of other finger-like protrusions such as microvilli, retraction fibers or nanotubes in different biological settings. As an example, FiloQuant successfully detected filopodia-like structures in activated natural killer cells. Although the role of these structures remain poorly documented in natural killer cells, similar cytoplasmic extensions have been described in T cells and suggested to contribute to T cell activation (Jung et al., 2016).

## Materials and methods

### FiloQuant installation

The files necessary to run FiloQuant in Fiji (https://fiji.sc/) are provided as supplemental software together with test images. Alternatively, FiloQuant installation in Fiji can be easily achieved through the ImageJ update site (see supplementary guide). Briefly, In Fiji, click on “Help → Update”, then “Manage update sites”, “add my site”. In the field “ImageJ Wiki account” input: “FiloQuant” then click “OK”. Close the “Manage update sites” window and, in the ImageJ Updater window, click on “Apply changes”. FiloQuant can then be found under “plugin → FiloQuant”. To run FiloQuant in ImageJ, users need to install the following dependencies: Enhanced Local Contrast (CLAHE.class;http://imagej.net/Enhance_Local_Contrast_(CLAHE)), Skeletonize3D.jar (http://imagej.net/Skeletonize3D), AnalyzeSkeleton.jar (http://imagej.net/AnalyzeSkeleton) (Arganda-Carreras et al., 2010) and Temporal-Color Code (http://imagej.net/Temporal-Color_Code).

### Cell culture and transient transfection

Immortalized normal breast epithelial cells (MCF10A), T24 c-Ha-ras oncogene-transfected MCF10A cells (MCF10AT) and invasive variant MCF10 DCIS.COM (DCIS.COM) cells were cultured in a 1:1 mix of Dulbecco’s Modified Eagle’s Medium (DMEM; Sigma-Aldrich) and F12 (Sigma-Aldrich) supplemented with 5% horse serum (Gibco, 16050-122), 20 ng/mL human epidermal growth factor (EGF; Sigma-Aldrich; E9644), 0.5 mg/ml hydrocortisone (Sigma-Aldrich, H0888-1G), 100 ng/ml cholera toxin (Sigma-Aldrich, C8052-1MG), 10 μg/ml insulin (Sigma-Aldrich, I9278-5ML) and 1% (v/v) penicillin/streptomycin (Sigma-Aldrich, P0781-100ML). DCIS.COM cells were cloned from a cell culture initiated from a xenograft obtained after two trocar passages of a lesion formed by MCF10AT cells (Miller et al., 2000). MCF10A Life-act and DCIS.COM Life-act cells were generated by lentiviral transduction (see method below). A2780 (ovarian carcinoma) cells were cultured in RPMI 1640 (Sigma-Aldrich) supplemented with 10% fetal calf serum (FCS). MDA-MB-231 (triple-negative human breast adenocarcinoma) cancer cells and telomerase immortalized human fibroblasts (TIFs) were grown in DMEM (Sigma-Aldrich) supplemented with 10% FCS. 293FT packaging cell line were grown in high glucose DMEM supplemented by 10% FCS, 0.1 mM non-essential amino acids, 1 mM Sodium Pyruvate, 6 mM L-glutamine, 1% (v/v) penicillin/streptomycin and 0.5 mg/ml geneticin (all from ThermoFisher Scientific). All cells were maintained at 37°C and 5% CO_2_.

Primary hippocampal neurons were isolated from E20 rat embryos. Briefly, embryonic brain tissue was dissected, and neurons were recovered by enzymatic digestion with trypsin and mechanical dissociation. Cells were maintained in neurobasal medium supplemented with 2% B27 supplement, 0.5 mM L-glutamine, 0.1 mg/mL primocin and 25 μM glutamate (all from Invitrogen).

NK-92 cells were maintained in alpha MEM complemented with 0.2 mM myoinositol, 0.1 mM beta-mercaptoethanol, 0.02 mM folic acid, 12.5% heat inactivated horse serum, 12.5% heat inactivated FCS (all from Sigma Aldrich), 2 mM L-glutamine and 1x non-essential amino acids (from Gibco). The growth medium was replaced every 2 days and supplemented with 100 U/ml of human recombinant interleukin-2 (Roche).

All cells were tested for mycoplasma contamination. Plasmids of interest were transfected using lipofectamine 3000 and the P3000TM Enhancer Reagent (Thermo Fisher Scientific) according to manufacturer’s instructions.

### Reagents, antibodies, plasmids and compounds

The anti-MAP2 antibody was acquired from Antibodies Online (ABIN372661; Aachen, Germany; used at 1:1000 for immunofluorescence). The Alexa Fluor^®^ 488 Phalloidin (A12379), used to stain filamentous actin, and DAPI (4′,6-diamidino-2-phenylindole, dihydrochloride, D1306) were purchased from Life Technologies. Bovine plasma fibronectin (FN) was purchased from Merck (341631). DMSO and latrunculin B (L5288-1MG) were obtained from Sigma-Aldrich. Growth factor reduced (GFR) Matrigel was bought from BD Biosciences (354230). *PureCol®* EZ Gel (fibrillar collagen I, concentration 5 mg/ml) was provided by Advanced BioMatrix. DQ™ Collagen (type I collagen from bovine Skin, fluorescein conjugate, D12060) was provided by Thermo Fisher Scientific. mEmerald-Lifeact-7 was a gift from Michael Davidson (Addgene plasmid # 54148). psPAX2 and pMD2.G were gifts from Didier Trono (Addgene plasmid # 12260 & # 12259). pCDH-Lifeact-mRFP was a gift from Dr Patrick Caswell (University of Manchester). Full-length bovine FN was labeled with Alexa Fluor^®^ 568 using an Alexa Fluor^®^ 568 Protein Labeling Kit (Thermo Fisher Scientific, A10238) according to manufacturer’s instructions.

### Virus production

Life-act mRFP lentiviral particles were generated in the 293FT packaging cell line following transient co-transfection of pCDH-Lifeact-mRFP, psPAX2 and pMD2.G constructs, in a 7:2:1 ratio, using the calcium-phosphate precipitation method (Graham and van der Eb, 1973). Virus-containing medium was collected 72 h post transfection, concentrated for 2 h at 25,000 rpm, resuspended in residual medium and flash frozen in liquid nitrogen. Functional titer was evaluated in 293FT cells by FACS (BD LSRFortessa, Becton Dickinson). To obtain stable Life-act expression, DCIS.COM cells were transduced with MOI 1 (multiplicity of infection; viral particle to cell number ratio of 1:1) and MOI 4 (viral particle to cell number ratio of 4:1) and MCF10A cells were transduced with MOI 4 and MOI 10 of viral stock. Cells exposed to different MOIs were then pooled 3 days post transduction and FACS sorted (BD FACSaria II cell sorter, Becton Dickinson, Franklin Lakes, US-NJ) with a gating strategy to obtain medium expression.

### Production of cell-derived matrices to monitor cell migration

Cell-derived matrices were generated as previously described (Jacquemet et al., 2013b). Briefly, TIFs were seeded at a density of 50,000 cells/ml in a 24-well plate. When confluent, cells were cultured for a further 10 days, with medium being changed every 48 h to complete medium supplemented with 50 μg/ml ascorbic acid (Sigma-Aldrich) to ensure collagen cross-linking. Mature matrices were then denuded of cells using lysis buffer (phosphate buffered saline (PBS) containing 20 mM NH4OH and 0.5 % (vol/vol) triton X-100). Following PBS washes, matrices were incubated with 10 mg/ml DNase I (Roche) at 37°C for 30 min. Matrices were then stored in PBS containing 1% (vol/vol) penicillin/streptomycin at 4°C prior to use.

### Circular invasion assay

A cartoon of the circular invasion assay protocol can be found in figure S1. Briefly, 5×10^4^ DCIS.COM or MCF10A cells were plated in one well of a culture-insert 2 well (ibidi) pre-inserted within a well of a μ-Slide 8 Well (ibidi). After 24 h, the culture-insert 2 well was removed and a gel of GFR Matrigel or fibrillar collagen (*PureCol^®^* EZ Gel) was casted. The gels were allowed to polymerise for 30 min at 37^°^C before normal media was added on top. Cells were left to invade for 3 days prior to fixation or live imaging (over 9 h).

### Proliferation assay

To monitor cell proliferation, cells were plated at low density in a well of a 6-well plate and imaged using a live-cell microscopy incubator (IncuCyte ZOOM^®^). Growth rates were calculated using the confluency method within the IncuCyte ZOOM software.

### 3D spheroid formation assay

To form spheroids in 3D Matrigel, cells were seeded as single cells, in normal growth media, at very low density (approximately 3000 cells per well) on GFR-Matrigel-coated glass-bottom dishes (MatTek, coverslip No. 0). After 12 h, the media was replaced by normal growth media supplemented with 2% (v/v) GFR Matrigel. The GFR Matrigel media was then changed every other day until the completion of the experiment.

### Zebrafish work

#### Zebrafish maintenance

Zebrafish (Danio rerio) housing and experimentation was performed under licence no. MMM/465/712-93 according to the European Convention for the Protection of Vertebrate Animals used for Experimental and other Scientific Purposes, and the Statutes 1076/85 and 62/2006 of The Animal Protection Law in Finland and EU Directive 86/609. Zebrafish were maintained and mated using standard procedures (Nüsslein-Volhard and Dahm, 2002; Westerfield, 2007).

#### Zebrafish intersegmental vessel sprouting assay

Transgenic zebrafish embryos expressing GFP in the endothelium (genotype Tg(kdrl:GFP); roy -/-; mitfa -/-) (Jin et al., 2005; White et al., 2008) were cultured at 28.5°C in E3-medium (5 mM NaCl, 0.17 mM KCl, 0.33 mM CaCl_2_, 0.33 mM MgSO_4_) prior to treatment with latrunculin B (150 ng/ml) or DMSO (1%) from 25 hours postfertilization (hpf) to 29 hpf. For live imaging of the sprouting segmental arteries, the embryos were dechorionated with forceps, anesthetized and mounted in 0.7% low-melting point agarose on glass-bottom dishes. Agarose was overlaid with E3 medium supplemented with tricaine (160 mg/ml, Sigma) and latrunculin B (150 ng/ml, Sigma) or DMSO (1%, Sigma). Imaging was performed at 28.5°C using a 3i spinning disk confocal microscope equipped with a 63x (NA 1.15) long distance water immersion objective.

#### Zebrafish embryo xenograft assay

Zebrafish embryos of the pigment-free casper strain (roy-/-; mitfa-/-) were used in the experiments. One 10 cm plate of MCF10 DCIS.COM cells stably expressing Life-act mRFP were trypsinized, washed twice in PBS and resuspended in 30 μl of 2% polyvinylpyrrolidone (Sigma) diluted in PBS for injection. Prior to injections, 24 hpf embryos were dechorionated, anaesthesized (160 mg/ml tricaine, Sigma) and immobilized with 0.7% low-melting point agarose (Sigma). Tumour cells were microinjected, using glass microinjection capillaries (TransferTip, Eppendorf), into the pericardiac cavity of 24 hpf zebrafish embryos using Celltram vario microinjector (Eppendorf) and Injectman (Eppendorf) micromanipulator mounted to SteroLumar V12 stereomicroscope (Zeiss). After injection, the embryos were released from the agarose with forceps, washed with E3 medium and cultured at 34°C in E3-medium. For imaging, the embryos were anaesthesized and mounted in low-melting point agarose on glass-bottom dishes.

### Microscopy set-up

The confocal microscope used was a laser scanning confocal microscope LSM780 (Carl Zeiss Microscopy, Thornwood, NY) with a 63x (NA 1.2 Water) objective controlled by the ZEN software (2010).

The spinning disk confocal microscope used was a Marianas spinning disk imaging system with a Yokogawa CSU-W1 scanning unit on an inverted Carl Zeiss Axio Observer Z1 microscope (Intelligent Imaging Innovations, Inc., Denver, USA). Objectives used were a 20x (NA 0.8 air, Plan Apochromat, DIC) objective (Carl Zeiss), a 63x oil (NA 1.4 Oil, Plan-Apochromat, M27 with DIC III Prism) objective (Carl Zeiss), a 63x water (NA 1.2 water C Apo, Korr C Apochromat UV-VIS-IR, DIC) objective (Carl Zeiss), a long working distance 63x water (NA 1.15 water, LD C-Apochromat, M27) objective or a 100x (NA 1.4 Oil, Plan-Apochromat, M27) objective. Images were acquired using either an Orca Flash 4 sCMOS camera (Chip size 2048x2048, Hamamatsu Photonics K.K., Hamamatsu City, Japan) or an Evolve 512 EMCCD camera (Chip size 512x512, Photometrics, Arizona, U.S).

The TIRF microscope used was a Zeiss Laser-TIRF 3 Imaging System (Carl Zeiss) equipped with a 100x (NA 1.46 Oil, alpha Plan-Apochromat, DIC) objective. Images were acquired on an EMCCD camera (Hamamatsu ImageEM C9100-13; Chip size 512x512; Hamamatsu Photonics K.K., Hamamatsu City, Japan) controlled by the Zen software (Zen 2012 Blue Edition Systems; Carl Zeiss).

The SIM-TIRF microscope used was an OMX SR (GE Healthcare Life Sciences, Issaquah, WA) fitted with a 60× Plan-Apo objective lens, 1.42 NA (immersion oil RI of 1.516) used in 2D-SIM-TIRF illumination mode (3 phases × 3 rotations within the TIRF plane per final image). Emitted light was collected on a front illuminated pco.edge sCMOS (PCO AG, Kelheim, Germany; pixel size 6.5 μm, readout speed 286 MHz) controlled by SoftWorx.

### Sample preparation for light microscopy

For TIRF microscopy experiments (related to figure 2B), cells transiently expressing bovine mCherry-Myosin-X were plated for 2 h on glass-bottom dishes (MatTek Corporation) precoated with 10 μg/ml of bovine plasma FN overnight at 4°C.

If not stated otherwise, all samples were fixed in 4% (wt/vol) paraformaldehyde (PFA) for 10 min, washed with PBS and permeabilized with PBS containing 0.5% (vol/vol) triton X-100 for 3 min. Cells were then washed with PBS, blocked using a solution of 1 M glycine for 30 min and incubated for 1 h at room temperature with Alexa Fluor^®^ 488 Phalloidin (1/100 in PBS) and, when indicated, with 1 μg/ml (in PBS) of dapi. After washing, samples were stored in PBS in the dark at 4^°^C prior to analysis. NK-92 natural killer cells were plated for 20 minutes on similar dishes precoated with 5 μg/ml of anti-CD18 antibody (clone IB4, produced in house) and 5 μg/ml of anti-NKp30 (Bio Legend, anti-human CD337, clone PG30.15).

If not stated otherwise, all live-cell imaging experiments were performed in normal growth media supplemented with 50 mM Hepes at 37°C and in the presence of 5% CO_2_.

### FiloQuant and TrackMate analysis of filopodia dynamics

To analyse filopodia dynamics in the circular invasion or 3D spheroid assays, filopodia were first detected and analyzed using the automated version of FiloQuant and the “stack analysis” option. In addition, filopodia further than 40 pixels away from the detected cell edge were excluded using the “maximal distance from cell edges” option. The tracking file generated by FiloQuant was then used as an input for TrackMate, an automated tracking software freely available within ImageJ (Tinevez et al., 2017). TrackMate was chosen, over other available ImageJ tracking plug-ins, because of its user-friendly interface and high flexibility. In TrackMate, the LoG detector (Estimated bob diameter = 0.1 μm; Threshold = 10; sub-pixel localization enabled) as well as the simple LAP tracker (Linking max distance = 1 μm; gap-closing max distance = 1 μm; Gap-closing max frame gap = 1) were used.

### Statistical analysis

The Tukey box plots represent the median and the 25th and 75th percentiles (interquartile range); points are displayed as outliers if 1.5 times above or below the interquartile range; outliers are represented by dots. Statistical analyses were performed when appropriate and P-values indicated by an asterisk in the figure legends. Unless otherwise indicated, the Student’s t-test was used (unpaired, two-tailed, unequal variance).

## Data availability and software updates

The authors declare that the data supporting the findings of this study are available within the article and from the authors on request. FiloQuant code is available as supplementary files associated with this article. Updates of FiloQuant will be released through the FiloQuant imageJ update site.

## Competing interests

The authors declare no competing interests.

## Author contributions

Conceptualization, G.J. and J.I.; methodology, G.J, I.P., A.F.C., A.P. and J.I.; formal analysis, G.J.; investigation, G.J., I.P., A.F.C, A.P.; resources, J.I. and J.S.O.; writing original draft, G.J. and H.H.; writing–reviewing, G.J., H.H., I.P., A.F.C., A.P., J.I; visualization, G.J.; supervision, G.J. and J.I.; funding acquisition, J.I. and J.S.O.

## Acknowledgement

We thank Hans-Juergen Kreienkamp (University Medical Center Hamburg-Eppendorf, Hamburg, Germany) for providing the primary hippocampal neuron samples. We thank Camilo Guzmán for his inputs in the software and the manuscript. We are grateful to Aki Stubb and Johanna Lilja for testing the FiloQuant software. We thank Patrick Caswell (University of Manchester) for providing reagents. The Cell Imaging Core and Zebrafish Core Facility (Turku Centre for Biotechnology, University of Turku and Åbo Akademi University) as well as the Live Cell Imaging Core Facility at the UT Southwestern Medical Center (Dallas, TX) are acknowledged for services, instrumentation and expertise.

## Funding

This study has been supported by the Academy of Finland (J.I.), an ERC Consolidator Grant (J.I.), the Sigrid Juselius Foundation (J.I.), the Finnish Cancer Organization (J.I.) and by the National Institute of Health (NIH-R01AI0679-11) (J.S.O). G.J. was supported by an EMBO Long-Term Fellowship.

---

**Figure S1:**
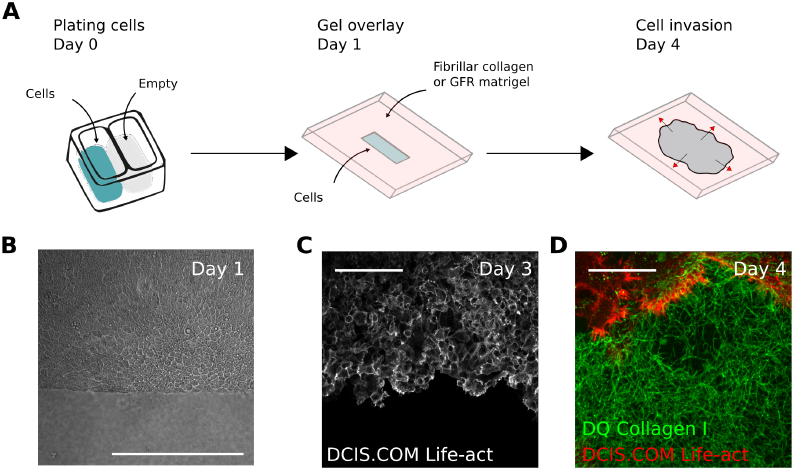
Circular invasion assay. **A:** Cartoon representing the different steps of the circular invasion protocol used in this study. Briefly, cells were seeded in a confined environment using a culture insert, the next day the insert was removed and a gel composed of either GFR Matrigel or fibrillar collagen I was casted and media added. Cells were then left to invade through the gel for 3 days prior to fixation or live imaging. **B:** Brightfield image of DCIS.COM cells after insert removal and gel overlay (day 1; scale bar = 400 μm). **C:** DCIS.COM Life-act cells were left to invade through fibrillar collagen I for 3 days and imaged using an SDC microscope (20x, Orca Flash 4 camera, scale bar = 100 μm). **D:** DCIS.COM Life-act cells were left to invade through fibrillar collagen spiked with 1μg.ml-1 of fluorescently labellebed collagen 1 prior to gel polymerisation (to visualize collagen architecture; DQ collagen) for 3 days and imaged live using an SDC microscope (100x, Evolve 512 EMCCD camera, scale bar = 25 μm).

**Video 1:** MCF10A and DCIS.COM cells stably expressing Life-act-mRFP were plated in circular invasion assays and left to invade through fibrillar collagen I or GFR Matrigel for 3 days before being imaged live on an SDC microscope (100x objective, Evolve 512 EMCCD camera) for over 9 h (1 picture every 3 min).

**Video 2:** FiloQuant analysis of a movie depicting DCIS.COM cell invasion through fibrillar collagen (circular invasion assay). The original movie (input), in addition to the multiple output files generated by FiloQuant, is displayed. Output files include filopodia detection (magenta), tracking image (filopodia dynamics), skeleton (mask used for filopodia detection), edge detection (defining the filopodia-free cell edge) and contour detection (to measure edge length).

**Video 3:** FiloQuant analysis of a movie depicting MCF10A cell invasion through fibrillar collagen (circular invasion assay). The original movie (input), in addition to a FiloQuant output of detected filopodia (magenta), is displayed.

**Video 4:** FiloQuant analysis of a movie depicting DCIS.COM cell invasion through fibrillar collagen (circular invasion assay). The original movie (input), in addition to a FiloQuant output of detected filopodia (magenta), is displayed.

**Video 5:** FiloQuant analysis of a movie depicting MCF10A cell invasion through GFR Matrigel (circular invasion assay). The original movie (input), in addition to a FiloQuant output of detected filopodia (magenta), is displayed.

**Video 6:** FiloQuant analysis of a movie of DCIS.COM cell invasion through GFR Matrigel. The original movie (input), in addition to a FiloQuant output of detected filopodia (magenta), is displayed.

**Video 7:** FiloQuant analysis of a movie monitoring a single DCIS.COM spheroid in 3D GFR Matrigel. DCIS.COM cells stably expressing Life-act-mRFP were plated in 3D GFR Matrigel for 3 days before being imaged live on an SDC microscope (100x objective, Evolve 512 EMCCD camera) for over 3 h (1 picture every 2 min). The original movie (input), in addition to the multiple output files generated by FiloQuant, is displayed. Output files include filopodia detection (magenta), tracking image (filopodia dynamics), skeleton (mask used for filopodia detection), edge detection (defining the filopodia-free cell edge) and contour detection (to measure edge length).

**Video 8:** Analysis of DCIS.COM spheroid rotation in 3D GFR Matrigel. DCIS.COM cells stably expressing Life-act-mRFP were plated in 3D GFR Matrigel for 3 days before being imaged live using an SDC microscope (100x objective, Evolve 512 EMCCD camera) for over 3 h (1 picture every 2 min).

**Video 9:** Analysis of DCIS.COM spheroid rotation in 3D GFR Matrigel. DCIS.COM cells stably expressing Life-act-mRFP were plated in 3D GFR Matrigel for 3 days before being imaged live using an SDC microscope (100x objective, Evolve 512 EMCCD camera) for over 3 h (1 picture every 2 min).

**Video 10:** Actin dynamics (single z plane) of a DCIS.COM spheroid growing in the pericardial cavity of a zebrafish embryo. DCIS.COM Life-act cells were injected in the pericardial cavity of zebrafish embryos and imaged live using an SDC microscope 24 h post-injection.

**Video 11:** Actin dynamics (3D reconstruction) of a DCIS.COM spheroid growing in the pericardial cavity of a zebrafish embryo. DCIS.COM Life-act cells were injected in the pericardial cavity of zebrafish embryos and imaged live using an SDC microscope 24 h post-injection. 3D reconstruction was generated using the Imaris software.

**Supplementary software 1**: Version of FiloQuant designed to analyse a single image already opened in ImageJ. This version of FiloQuant contains step-by-step user validation of the various processing stages to help users achieve optimal settings for filopodia detection.

**Supplementary software 2**: Version of FiloQuant designed to analyse images within a specified folder. This version of FiloQuant still contains step-by-step user validation of the various processing stages to help users achieve optimal settings for filopodia detection.

**Supplementary software 3:** Version of FiloQuant designed to automatically analyse mages within a specified folder by using the same settings for all images (batch analyses). This version of FiloQuant also allows the analysis of stacks / movies and provides a tracking file as well as a time projection of the detected filopodia when the “stack analysis” option is enabled.

